# Glutamatergic Systems in Ctenophores

**DOI:** 10.64898/2026.08.09.743775

**Authors:** Leonid L. Moroz, Tigran P. Norekian

**Author notes:** **Corresponding author:** Leonid L. Moroz.

## Abstract

Despite glutamate’s widespread role as the dominant excitatory transmitter in vertebrate brains, the early evolution of glutamate and its recruitment into neural signaling remain largely unknown. The major limitation is the lack of information on its distribution in early-branching basal metazoans, such as ctenophores (comb jellies). Here, using glutamate immunoreactivity (IR) in two ctenophore species with distinct ecologies (*Pleurobrachia bachei* and *Beroe abyssicola*), we show that glutamate IR is present in subpopulations of neurons within the subepithelial neural network and in small groups of mesogleal neuron-like cells, and that it differentially labels some muscle fibers. Remarkably, we also observed an enriched glutamate-ir signal within the nuclei of subepithelial neurons in *Beroe*. However, glutamate expression levels are species-specific, suggesting a tight coupling of glutamate recruitment for neural communication with energetic demands.

## Introduction

L-glutamate is one of the most abundant, versatile, and still enigmatic structural and regulatory metabolites, directly involved in more than 200 biochemical pathways and in countless integrative systems as a signaling molecule across all domains of Life, from prokaryotes to eukaryotes (Moroz et al., 2021a). In vertebrate brains, glutamatergic synapses mediate 50-80% of interneuronal communications, where glutamate is an excitatory neurotransmitter (Micheva et al., 2010), primarily in interneurons and some sensory cells. However, in studied invertebrates, glutamatergic neurons (i.e., neurons that release and use glutamate as a transmitter) represent a relatively small fraction of neuronal populations - usually less than 10% (Moroz et al., 2021a). Notably, across phyla, glutamate acts as both an inhibitory (Kehoe et al., 2009) and an excitatory transmitter; these effects are mediated by different classes of ionotropic (iGluRs), pentameric cys-loop receptor channels (Cymes and Grosman, 2021; Jaiteh et al., 2016; Kehoe et al., 2009; Lynagh et al., 2015) and metabotropic receptors, most of which were lost in the lineage that led to vertebrates (Moroz et al., 2021a).

It was proposed that the complexity of ionotropic glutamate receptors (iGluRs) correlates with the establishment of multicellularity; metazoans possess one of the greatest diversities of iGluRs (Moroz et al., 2021a). However, the early evolution of animal glutamate signaling and its recruitment into neural circuits remain enigmatic. To date, no glutamatergic neurons have been reliably identified or even mapped in representatives of prebilaterian metazoans such as ctenophores. Ctenophores, or comb jellies, the lineage sister to the rest of animals, independently evolved neural systems with the most distinct arrays of transmitters (Moroz et al., 2014, Moroz, 2015, Moroz, Kohn, 2016). In addition to several dozen ctenophore-specific (neuro)peptides, the only proposed small-molecule transmitter was glutamate as a neuromuscular messenger (Moroz et al., 2014). This hypothesis, based on observations in *Pleurobrachia bachei* as the major model, held that both L- and D-glutamate induced action potentials in specific muscle cells, increased intracellular Ca^2+^, and induced muscle contractions. Notably, D-glutamate was also detected in ctenophores but not in other studied metazoans (Moroz et al., 2020); however, L-glutamate is the most abundant and the most potent in inducing muscular contractions (Moroz et al., 2014).

Genomic, transcriptomic, and in situ hybridization data revealed 10-15 epsilon-type iGluRs (Moroz et al., 2014, 2021a). Cloning and expression of some of these receptors in *Xenopus* oocytes showed that some are sensitive to glycine (Alberstein et al., 2015; Yu et al., 2016). Furthermore, no vesicular glutamate transporter was identified in ctenophores (Moroz et al., 2021a; Moroz, Kohn, 2015); instead, we found a subclass of ctenophore-specific sialin-like (SLC17-type) transporters (Moroz et al., 2021a). Mapping glutamatergic neurons is challenging because in situ hybridization with most markers labels cell somata rather than processes and synaptic areas, as most mRNAs are not transported there or are not detectable due to low abundance. Thus, it is difficult to reliably visualize glutamatergic neurons and distinguish them from other secretory cells.

Here, we used glutamate-specific antibodies to characterize the distribution of putative glutamatergic neurons in representatives of two reference genera with sequenced genomes: *Pleurobrachia bachei* and *Beroe abyssicola* (Moroz et al., 2014; Vargas et al., 2024), revealing neuron- and muscle-specific patterns of Glu IR across epithelial and mesodermal cell populations. We selected these species for their abundance and markedly different ecologies. *Pleurobrachia* is an ambush predator with a pair of tentacles that captures various crustaceans in zooplankton, whereas *Beroe* is an active predator without tentacles that hunts other ctenophores and swallows them with a large mouth and cilia-based teeth (Tamm, 2014). Equally important for our selection was that these species are relatively easy to fix and suitable for immunohistochemical studies of adult animals (Norekian, Moroz, 2016, 2019, 2021). In contrast, most lobates, including popular *Mnemiopsis* and *Bolinopsis* species, are extremely fragile; it is virtually impossible to fix and work on adults.

**Figure 1.**
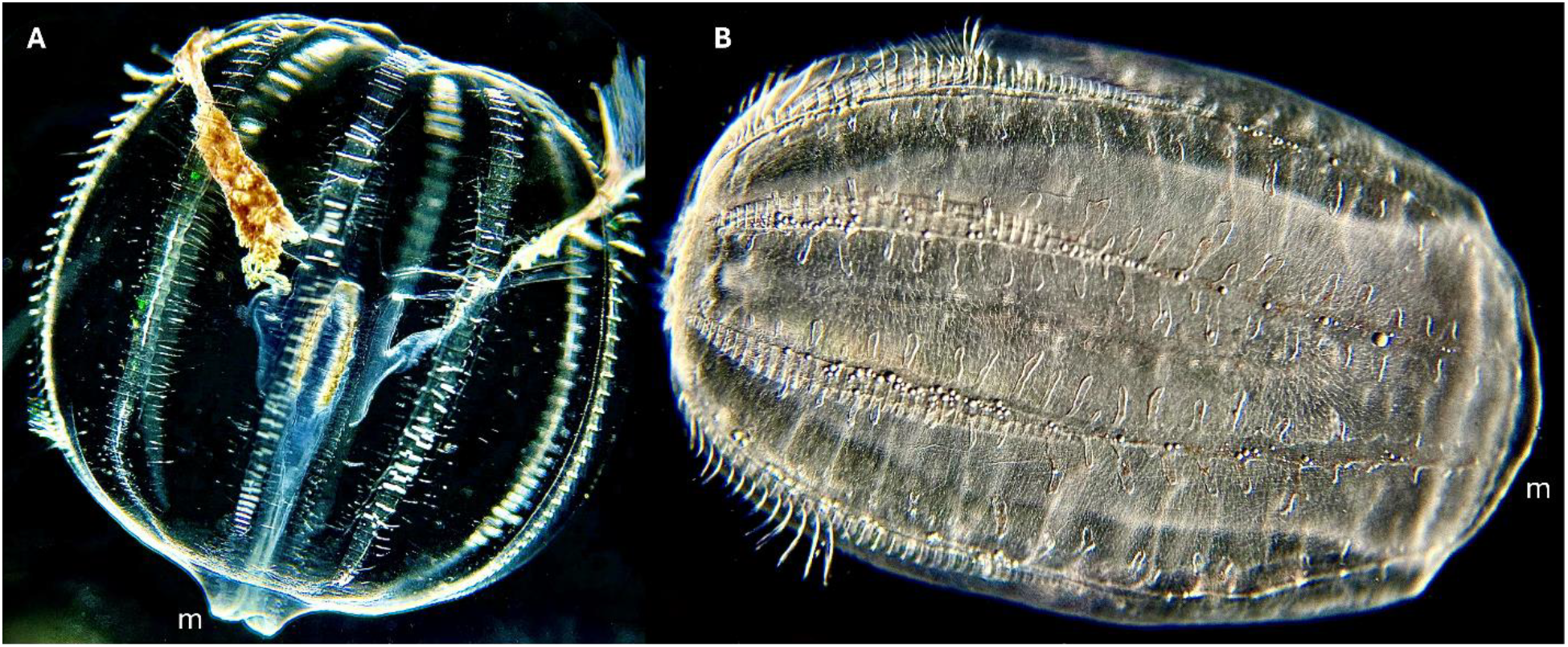
**Ctenophores used in the current study**: *Pleurobrachia bachei* (A) and *Beroe abyssicola* (B). These are transparent pelagic comb jellies with ciliated locomotion mediated by eight comb rows. *Pleurobrachia* is an ambush predator that feeds mostly on crustaceans, whereas *Beroe* feeds on other ctenophores, primarily *Bolinopsis* and occasionally on *Pleurobrachia*.

## Materials and Methods

### Animals

Adult specimens of *Pleurobrachia bachei* and *Beroe abyssicola* were collected from the dock in the North Pacific Ocean at Friday Harbor Laboratories (University of Washington) and held in 1-gallon jars at 10° C.

### Immunocytochemistry and Phalloidin staining

Adult animals were fixed overnight (10-12 hours) in 4% paraformaldehyde in 0.1 M phosphate-buffered saline (PBS) at +5º C and washed for 2 hours in PBS. The dissected tissues of fixed adult animals were pre-incubated overnight in a blocking solution of 6% goat serum in PBS containing 0.02% Triton X-100 (PBT). The samples were then incubated for 48 hours at +5 °C in primary antibodies diluted in 6% goat serum to a final dilution of 1:200. We used polyclonal anti-glutamate antibodies produced in rabbit (Sigma Cat#G6642, RRID: AB_259946). Following a series of PBS washes for 6-8 hours, the dissected tissues of adult animals were incubated for 20 hours in secondary goat anti-rabbit IgG antibodies: conjugated to Alexa Fluor 568 (ThermoFisher Cat# A-11011, RRID: AB_143157) at a final dilution of 1:100.

We also used rat monoclonal antibodies against tubulin (AbD Serotec Cat# MCA77G, RRID: AB_325003) in double-labeling experiments at a final dilution of 1:100 to identify overall populations of neurons (see Norekian and Moroz 2019a, b). This type of antibodies recognizes the alpha subunit of tubulin and specifically bind tyrosylated tubulin (Wehland & Willingham, 1983; Wehland et al., 1983). After a series of PBS washes for 6-8 hours, the dissected tissues were incubated for 12 hours in secondary goat anti-rat IgG antibodies: Alexa Fluor 488 conjugated (Molecular Probes, Invitrogen, Cat# A11006, RRID: AB_141373) at a final dilution of 1:100.

To label the muscle fibers, we used the well-known marker phalloidin (Alexa Fluor 488 phalloidin from Molecular Probes), which binds to F-actin (Wulf, Deboben, Bautz, Faulstich, & Wieland, 1979). After washing in PBS following the secondary antibody treatment, the samples were incubated in phalloidin solution (in PBS) for 6 to 8 hours at +5º C, at a final dilution of 1:100, and then washed in several PBS rinses for 8 hours.

To stain the nuclei, dissected adult tissues were mounted in Antifade Mounting Medium with DAPI (Vectashield; Cat#H-2000). Slides were examined on a Nikon Research Microscope Eclipse E800 with epifluorescence using standard TRITC and FITC filters, and images were recorded on a Nikon C1 laser scanning confocal microscope. Total number of animals used in this study: *Beroe* – 16 adult animals; *Pleurobrachia* – 14 adult animals (all dissected into several smaller pieces for processing).

## Results

Both classical (Hernandez-Nicaise, 1991; Moroz, 2015) and more recent immunohistochemical (Norekian, Moroz, 2016, 2019, 2021), and volume microscopy (Burkhardt et al., 2023, Jokura et al., 2026; Ferraioli et al., 2026) data identified two distinct neural systems in ctenophores. These are the subepithelial polygonal net in the skin and a mesh of neuron-like cells in the mesoglea. We detected cell-specific glutamate immunoreactivity (Glu IR) in both the subepithelial net and the mesogleal region of *Beroe* and *Pleurobrachia*. However, using identical protocols and procedures, Glu-ir labeling in the polygonal net of *Beroe* was much more pronounced and clearer than in *Pleurobrachia* (not shown). Notably, only a fraction of neurons and their processes in *Beroe* polygonal net were labeled with glutamate IR, while some parts of the network identified by tubulin IR were not glutamate immunoreactive (glutamate-ir, Fig. 2). These glutamate-ir neurons were predominantly bipolar or tripolar and mostly located at the nodes of the net (Fig. 2, 3). In all instances, glutamate IR was co-localized with tubulin IR. However, the intracellular distribution of immunolabeling was markedly different. Tubulin IR was predominantly located in the neuronal processes, whereas glutamate IR was very strong in the cell somata (Fig. 2B). Interestingly, we detected the highest intensity of glutamate IR labeling in the neuronal nuclei (Fig. 2B, Fig. 3). No glutamate IR was observed in other types of cells in the subepithelial layer.

**Figure 2.**
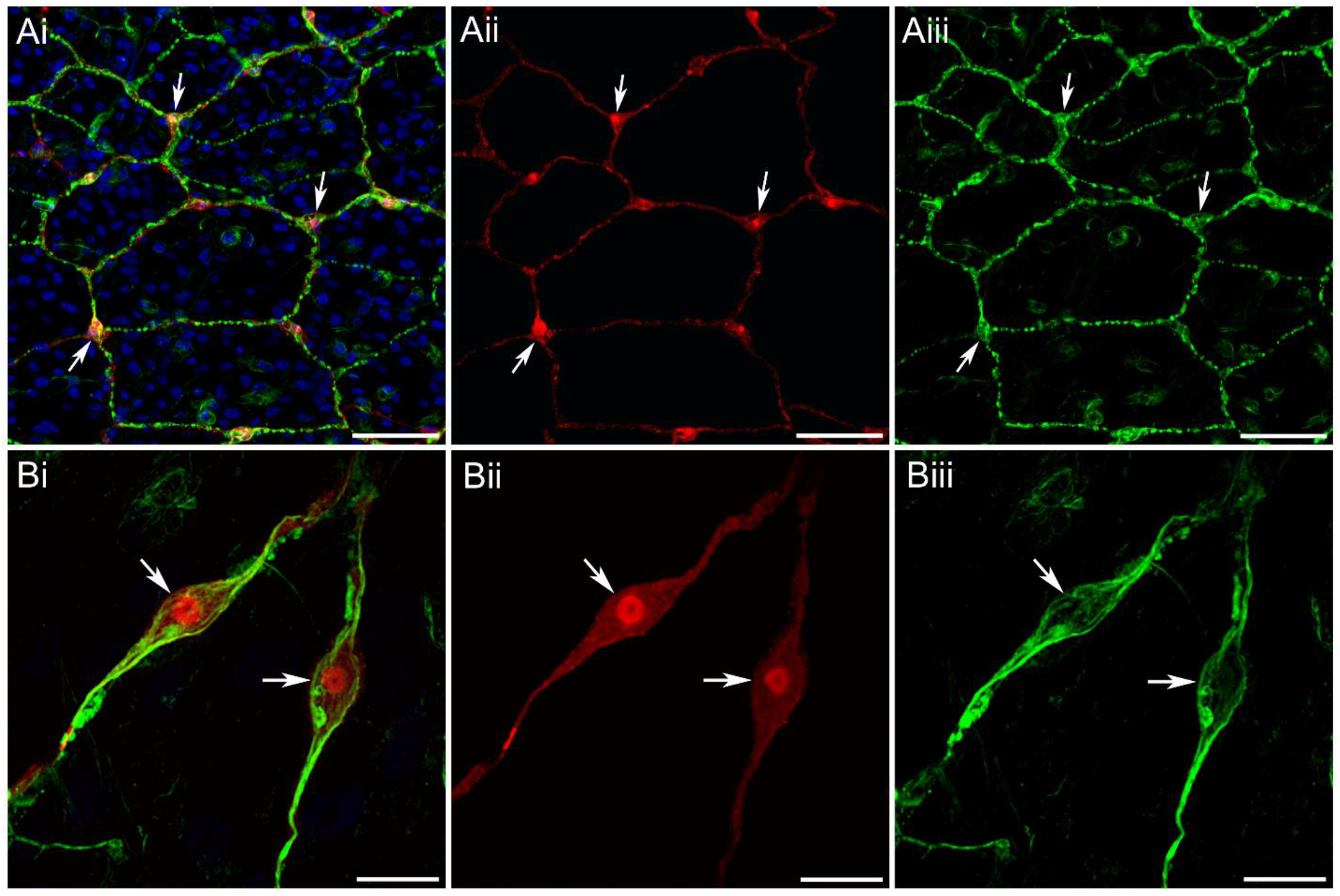
Double-labeling of the subepithelial polygonal neural net in *Beroe* with tubulin antibodies (green) and Glutamate antibodies (red). DAPI is blue. Ai – Polygonal neural net labeled with tubulin IR also contains numerous neurons, which are double-labeled with Glutamate IR (some are shown by arrows). However, not all elements of the tubulin-IR network are double-labeled with Glutamate IR. Aii – Red channel with Glutamate IR only. Aiii – Green channel with tubulin IR only. Bi – Two bipolar neurons from the polygonal network double-labeled with tubulin IR and Glutamate IR. Note that Glutamate antibodies predominantly label neural cell bodies, while tubulin antibodies better stain neural processes (see also Fig. 1B). Bii – Red channel with Glutamate IR only. Biii – Green channel with tubulin IR only. Scale bars: A – 50 µm, B – 10 µm.

**Figure 3.**
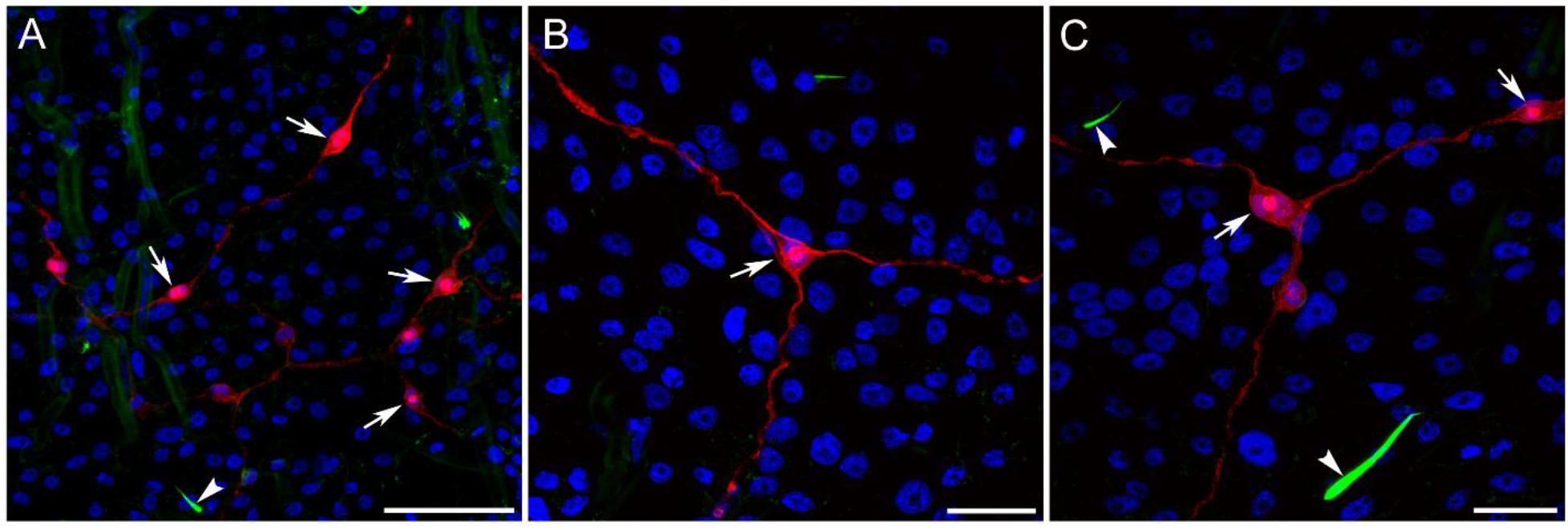
Glutamate-ir neurons in the subepithelial layer of the *Beroe abyssicola* skin. Glutamate IR is red, phalloidin is green, and DAPI is blue. A – Lower magnification shows a scattered distribution of putative glutaminergic neurons. B, C – Higher magnification shows individual neurons. Many neurons are bipolar. Arrows point to the neuronal cell bodies. Note the intense nuclear labeling in each stained neuron, suggesting a higher concentration of glutamate in nuclear compartments. Arrowheads indicate the surface sensory cilia of other neural types labeled with phalloidin. Scale bars: A – 50 µm, B, C – 20 µm.

The meshwork of glutamate-ir mesogleal cells was more prominent in *Pleurobrachia* (Fig. 4) than in *Beroe*. These cells exhibited diverse morphology, ranging from elongated bipolar to tripolar and multipolar, neuron-like cells with thin, extending processes (Fig. 4). Not all mesogleal cells identified by tubulin IR were glutamate-ir; only about half showed double-labeling (Fig. 4A). We did not detect differential nuclear labeling in these cells, unlike glutamate-ir neurons from the Beroe polygonal network, although cell somata were always brightly stained (Fig. 4B-D).

**Figure 4.**
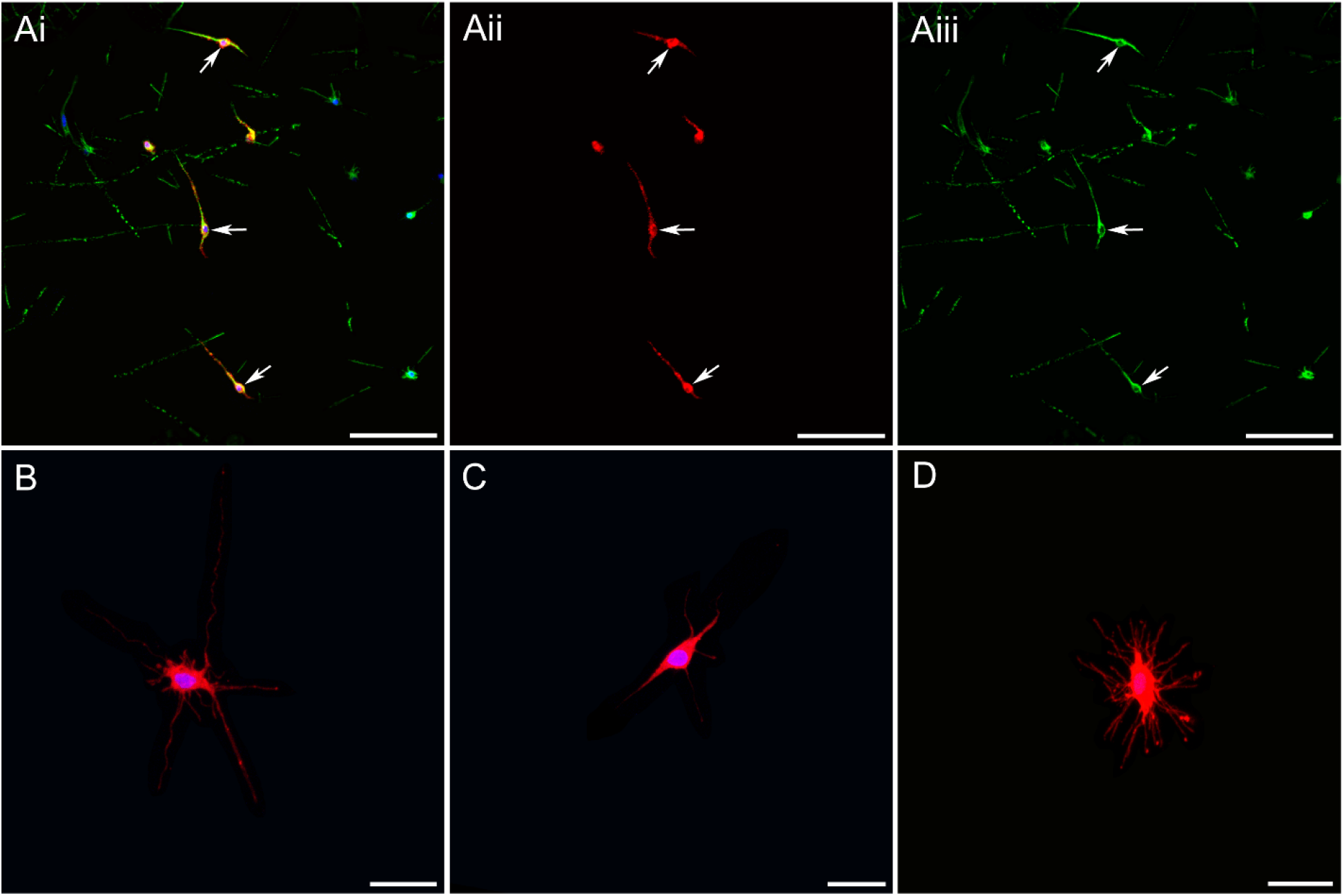
Mesogleal neuron-like cells in *Pleurobrachia*. Tubulin IR is green, Glutamate IR is red, while DAPI is blue. Ai – Some mesogleal neural cells, which are always stained with tubulin antibodies, are double-labeled with Glutamate antibodies (shown by arrows). Aii – Red channel with Glutamate IR only. Aiii – Green channel with tubulin IR only. B, C, D – Morphological structure of different Glutamate-ir mesogleal neural cells (no tubulin IR). Scale bars: A – 50 µm, B – 10 µm, C – 10 µm, D – 10 µm.

**Figure 5.**
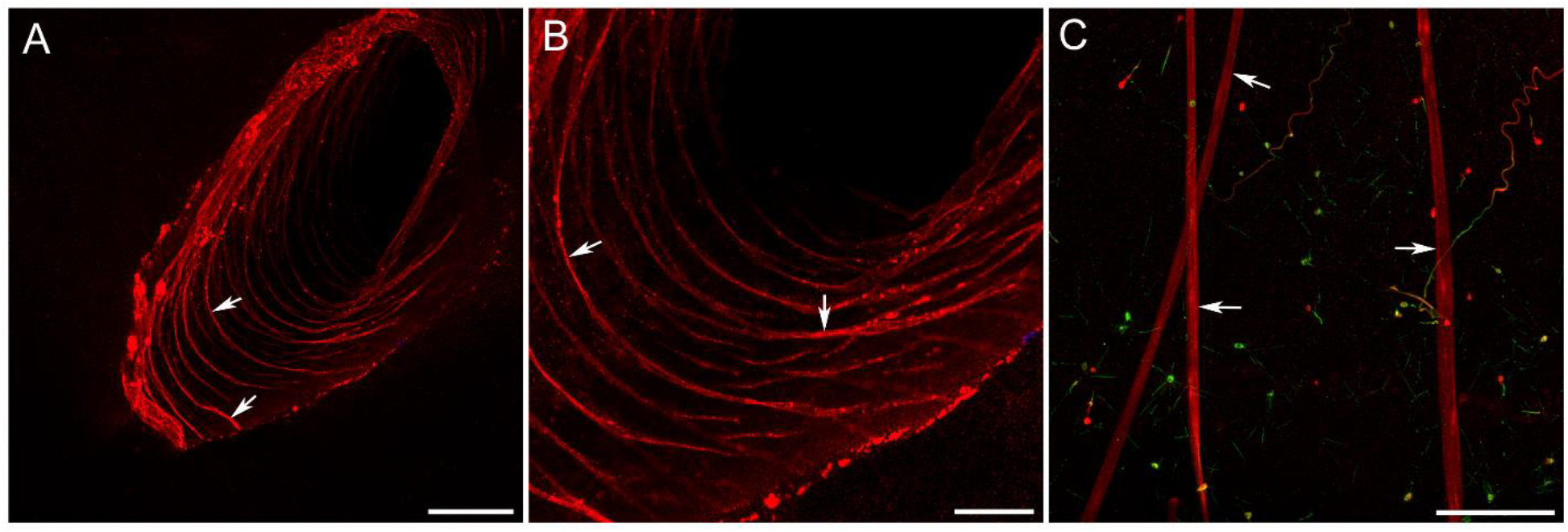
Muscle fibers in *Pleurobrachia* are labeled with Glutamate antibodies (red). A, B – Circular muscle fibers in the wall of the tentacle pocket (some are shown by arrows). C – Large muscle fibers (arrows) in the mesoglea region, also labeled with Glutamate IR. Tubulin IR is green. Scale bars: A – 100 µm, B – 50 µm, C – 100 µm.

In addition to various neuronal cells, glutamate IR also labeled different muscle fibers in *Pleurobrachia*. We identified a population of glutamate-ir circular muscles within the wall of a tentacle pocket, spanning its entire depth (5A, B). Very long, large-diameter muscle fibers crossing the mesoglea region also showed strong Glu-ir labeling, suggesting a predominant accumulation of glutamate in these contractile cell types. Such strong labeling was not observed in *Beroe*, where we saw only a very weak signal from some muscle fibers, implying different metabolic constraints for muscle types across ctenophores.

## Discussion

The observed differential neuronal immunolabeling within the polygonal net is consistent with the hypothesis that glutamate can serve as a neuromuscular transmitter (Moroz et al., 2014). Subepithelial muscles were used in prior electrophysiological, patch-clamp, and imaging experiments, which showed their sensitivity to glutamate but not to other canonical small-molecular-weight transmitters tested (acetylcholine, histamine, dopamine, noradrenaline, adrenaline, octopamine, serotonin, GABA, nitric oxide). It was also shown that some processes of neurons from the subepithelial net make direct contact with the underlying muscles, although synapses have not yet been identified, suggesting that glutamate can also act via volume transmission, at least in certain cases.

We also anticipate that some populations of subepithelial neurons in ctenophores do not use glutamate and might be purely peptidergic (Moroz, et al., 2014; Sachkova et al., 2021, Hayakawa et al., 2022) or use other transmitter classes, potentially with multiple targets. In larval and early juvenile *Mnemiopsis*, at least some parts of the subepithelial nets can form cytoplasmic bridges and may constitute syncytial nets (Burkhardt et al., 2023, Jokura et al., 2026; Ferraioli et al., 2026). However, this has not been confirmed in adult animals or in the species used in the current studies. In addition, each cell in the net contains secretory vesicles and may participate in chemical transmission via localized or volume signaling (Moroz, 2023), which is the most ancestral form of intraneuronal communication in early metazoans without synapses (Moroz, 2009; 2014, 2021; Moroz et al., 2021b; Jekely, 2021). The functional predominance of chemical transmission is supported by experiments with high Mg^*2+*^*-*Ca^*2+*^ that suppress secretion and behavioral integration (Norekian, Moroz, 2024). At present, we do not know the relative contributions of different (neuro)transmitters to behavioral integration.

This study illustrates that the scope and patterns of Glu IR are both species- and cell-specific, suggesting a tight coupling of glutamate recruitment to ecology across distributed integrative, feeding, and locomotory systems with varying bioenergetic demands. Nevertheless, outcomes of glutamate recruitment cannot be predicted at this stage of knowledge. In retrospect, we can propose that the more prominent labeling in *Beroe* neurons might reflect the greater neural coordination required by these ferocious predators compared with *Pleurobrachia*, but larger neuron sizes should also be considered. In contrast, the more intense labeling of mesogleal neural-like cells in *Pleurobrachia* is less explainable at this point; *Beroe* has relatively smaller fractions of these cells embedded in highly developed muscular systems with truly giant muscular cells.

Muscle labeling in *Pleurobrachia* is more intense than in *Beroe*; this may be associated with the maintenance of tonic activity, as a hypothesis. Glutamate functions in ctenophore muscles can be both signaling and metabolic. Glutamate is the most dynamic and vital among anaplerotic substances (i.e., molecules responsible for refilling catabolic intermediates of the TCA cycle (Brunengraber, H., Roe, C. R., 2006), as illustrated by a decline in intramuscular glutamate at the start of exercise. By deamination into α-ketoglutarate, glutamate is the principal energy substrate fueling the TCA cycle, both in muscles and the brain (Krebs, 1953; Bordone et al., 2019, McKenna, 2013). The net energy yield from the oxidation of one molecule of glutamate is 23-26 ATP molecules (McKenna, 2013). It might be rational to repeatedly recruit Glu in cell types and cell states with high energy demands.

Furthermore, in ctenophores, as in other animal clades, we expect biochemical interactions between glutamate metabolism and tubulin because glutamination of tubulin (Bigman, Levy, 2020; Edde et al., 1990) is a well-known mechanism that regulates cytoskeletal dynamics, including motile (Kubo, Oda, 2017; Wang et al., 2021) and non-motile, sensory cilia (Kimura et al., 2018; Yang et al., 2021), axonal transport (Kubo et al., 2010), and synapse functions (Akella, Barr, 2021; Ikegami et al., 2007).

Finally, one unexpected finding of this work is the localization of glutamate in the nuclei of *Beroe* neurons. Due to its distinct properties, potassium glutamate acts as an osmolyte and chaperone that stabilizes protein complexes (Young, Ajami, 2000; Cheng et al., 2016) and protein-DNA interactions (Deredge et al., 2010; Leirmo et al., 1987; Sengupta et al., 2016), including nucleosome assembly (Regnard et al., 2000) and activity of many DNA-modifying enzymes (Hanish, McClelland, 1988). In addition, many chromatin modifications are energy-demanding (although it is unclear whether the TCA operates in the ctenophore nuclei).

In sum, we are at the very beginning of exploring the roles of glutamate signaling in ctenophores, and the presented localization studies provide insights into its versatile functionality, which likely dates back to the dawn of animal multicellularity and even the dawn of life itself (Moroz et al., 2021a). In this perspective, ctenophores, as descendants of the earliest metazoan lineage, will provide unexpected insights into the transmitter and integrative functions of glutamate, one of the first intercellular signal molecules.

### Conclusion, limitations and future directions

The role of glutamate as a small-molecule neurotransmitter candidate is supported by prior pharmacological and electrophysiological experiments, as well as by its enrichment in subepithelial nerve nets. Nevertheless, physiological, presumably neuron-specific glutamate release needs to be studied alongside its intracellular localization. Whether glutamate is stored in secretory vesicles and the mechanisms of its uptake remain unclear, since no canonical vesicular glutamate transporter has been identified in sequenced ctenophore genomes. Equally important is the characterization of glutamate co-transmission with secretory peptides and the elucidation of the systemic behavioral functions of glutamate via synaptic messenger and/or volume transmission, which can also be non-neuronal (e.g., release from muscles and other cell types). The presence, localization, and role of D-glutamate also warrant further investigation.

Notably, glutamate is a key molecule in many metabolic pathways, and because it is one of the most abundant metabolites, high concentrations of glutamate can be associated with injury signaling in first metazoans via paracrine signaling (Moroz, 2009). This might well be an ancestral exaptation, explaining why glutamate was repeatedly recruited as a neurotransmitter in early animals in general and in ctenophores in particular. Finally, the enigmatic, potentially integrative functions of glutamate in neuronal nuclei could be a new, exciting direction for exploratory research.

In any of these endeavors, we advocate broader comparative analysis that encompasses the extant diversity of ctenophores, from pelagic to benthic species (Moroz et al., 2024). This approach would likely be the most informative, given the observed differences between just two species studied, which serve as unique natural experiments with unpredictable ‘solutions’ to environmental stress and complex adaptations.

## Acknowledgments

We thank FHL for their excellent facilities, including the Nikon Laser Scanning confocal microscope. This research was supported by the National Science Foundation grant (IOS-2341882) and National Institute of Health grant (5R01NS11449) to LLM.

## Data Availability Statement

The data that support the findings of this study are available from the corresponding author upon reasonable request.

## References

Akella, J. S., Barr, M. M., 2021. The tubulin code specializes neuronal cilia for extracellular vesicle release. Dev Neurobiol 81, 231–252.

Alberstein, R., Grey, R., Zimmet, A., Simmons, D. K., Mayer, M. L., 2015. Glycine activated ion channel subunits encoded by ctenophore glutamate receptor genes. Proc Natl Acad Sci U S A 112, E6048-6057.

Bigman, L. S., Levy, Y., 2020. Tubulin tails and their modifications regulate protein diffusion on microtubules. Proc Natl Acad Sci U S A 117, 8876–8883.

Bordone, M. P., Salman, M. M., Titus, H. E., Amini, E., Andersen, J. V., Chakraborti, B., Diuba, A. V., Dubouskaya, T. G., Ehrke, E., Espindola de Freitas, A., Braga de Freitas, G., Goncalves, R. A., Gupta, D., Gupta, R., Ha, S. R., Hemming, I. A., Jaggar, M., Jakobsen, E., Kumari, P., Lakkappa, N., Marsh, A. P. L., Mitlohner, J., Ogawa, Y., Paidi, R. K., Ribeiro, F. C., Salamian, A., Saleem, S., Sharma, S., Silva, J. M., Singh, S., Sulakhiya, K., Tefera, T. W., Vafadari, B., Yadav, A., Yamazaki, R., Seidenbecher, C. I., 2019. The energetic brain - A review from students to students. J Neurochem 151, 139–165.

Burkhardt P, Colgren J, Medhus A, Digel L, Naumann B, Soto-Angel JJ, Nordmann EL, Sachkova MY, Kittelmann M. 2023. Science. Syncytial nerve net in a ctenophore adds insights on the evolution of nervous systems. 380(6642):293–297. doi: 10.1126/science.ade5645.

Brunengraber, H., Roe, C. R., 2006. Anaplerotic molecules: current and future. J Inherit Metab Dis 29, 327–331.

Cheng, X., Guinn, E. J., Buechel, E., Wong, R., Sengupta, R., Shkel, I. A., Record, M. T., Jr., 2016. Basis of Protein Stabilization by K Glutamate: Unfavorable Interactions with Carbon, Oxygen Groups. Biophys J 111, 1854–1865.

Cymes, G. D., Grosman, C., 2021. Signal transduction through Cys-loop receptors is mediated by the nonspecific bumping of closely apposed domains. Proc Natl Acad Sci U S A 118.

Deredge, D. J., Baker, J. T., Datta, K., Licata, V. J., 2010. The glutamate effect on DNA binding by pol I DNA polymerases: osmotic stress and the effective reversal of salt linkage. J Mol Biol 401, 223–238.

Edde, B., Rossier, J., Le Caer, J. P., Desbruyeres, E., Gros, F., Denoulet, P., 1990. Posttranslational glutamylation of alpha-tubulin. Science 247, 83–85.

Ferraioli A, Digel L, Sturm D, Colgren J, Le Goff C, Jan A, Soto-Angel JJ, Naumann B, Kittelmann M, Burkhardt P. 2026. The 3D architecture of the ctenophore aboral organ and the evolution of complex integrative centers in animals. Sci Adv. 2026 Mar 6;12(10):eaea8399. doi: 10.1126/sciadv.aea8399.

Hanish, J., McClelland, M., 1988. Activity of DNA modification and restriction enzymes in KGB, a potassium glutamate buffer. Gene Anal Tech 5, 105–107.

Hayakawa E, Guzman C, Horiguchi O, Kawano C, Shiraishi A, Mohri K, Lin MF, Nakamura R, Nakamura R, Kawai E, Komoto S, Jokura K, Shiba K, Shigenobu S, Satake H, Inaba K, Watanabe H. 2022 Nat Ecol Evol. Mass spectrometry of short peptides reveals common features of metazoan peptidergic neurons. Oct;6(10):1438–1448. doi: 10.1038/s41559-022-01835-7.

Hernandez-Nicaise, M.-L. (1991). “Ctenophora” in Microscopic Anatomy of Invertebrates: Placozoa, Porifera, Cnidaria, and Ctenophora. eds. Harrison, F. W. F. W., Westfall, J. A. (New York: Wiley), 359–418.

Ikegami, K., Heier, R. L., Taruishi, M., Takagi, H., Mukai, M., Shimma, S., Taira, S., Hatanaka, K., Morone, N., Yao, I., Campbell, P. K., Yuasa, S., Janke, C., Macgregor, G. R., Setou, M., 2007. Loss of alpha-tubulin polyglutamylation in ROSA22 mice is associated with abnormal targeting of KIF1A and modulated synaptic function. Proc Natl Acad Sci U S A 104, 3213–3218.

Jaiteh, M., Taly, A., Henin, J., 2016. Evolution of Pentameric Ligand-Gated Ion Channels: Pro-Loop Receptors. PLoS One 11, e0151934.

Jékely G. 2021. The chemical brain hypothesis for the origin of nervous systems. Philos Trans R Soc Lond B Biol Sci. Mar 29;376(1821):20190761. doi: 10.1098/rstb.2019.0761.

Jokura, K., Jasek, S., Niederhaus, L., Burkhardt, P., Jékely, G. 2026. Elife. Neural connectome of the ctenophore statocyst.14:RP108420. doi: 10.7554/eLife.108420.

Kehoe, J., Buldakova, S., Acher, F., Dent, J., Bregestovski, P., Bradley, J., 2009. Aplysia cys-loop glutamate-gated chloride channels reveal convergent evolution of ligand specificity. J Mol Evol 69, 125–141.

Kimura, Y., Tsutsumi, K., Konno, A., Ikegami, K., Hameed, S., Kaneko, T., Kaplan, O. I., Teramoto, T., Fujiwara, M., Ishihara, T., Blacque, O. E., Setou, M., 2018. Environmental responsiveness of tubulin glutamylation in sensory cilia is regulated by the p38 MAPK pathway. Sci Rep 8, 8392.

Kubo, T., Oda, T., 2017. Electrostatic interaction between polyglutamylated tubulin and the nexin-dynein regulatory complex regulates flagellar motility. Mol Biol Cell 28, 2260–2266.

Kubo, T., Yanagisawa, H. A., Yagi, T., Hirono, M., Kamiya, R., 2010. Tubulin polyglutamylation regulates axonemal motility by modulating activities of inner-arm dyneins. Curr Biol 20, 441–445.

Leirmo, S., Harrison, C., Cayley, D. S., Burgess, R. R., Record, M. T., Jr., 1987. Replacement of potassium chloride by potassium glutamate dramatically enhances protein-DNA interactions in vitro. Biochemistry 26, 2095–2101.

Lynagh, T., Beech, R. N., Lalande, M. J., Keller, K., Cromer, B. A., Wolstenholme, A. J., Laube, B., 2015. Molecular basis for convergent evolution of glutamate recognition by pentameric ligand-gated ion channels. Sci Rep 5, 8558.

McKenna, M. C., 2013. Glutamate pays its own way in astrocytes. Front Endocrinol (Lausanne) 4, 191.

McLaggan, D., Naprstek, J., Buurman, E. T., Epstein, W., 1994. Interdependence of K+ and glutamate accumulation during osmotic adaptation of Escherichia coli. J Biol Chem 269, 1911–1917.

Micheva, K. D., Busse, B., Weiler, N. C., O’Rourke, N., Smith, S. J., 2010. Single-synapse analysis of a diverse synapse population: proteomic imaging methods and markers. Neuron 68, 639–653.

Moroz, L. L., 2009. On the independent origins of complex brains and neurons. Brain Behav Evol 74, 177–190. doi: 10.1159/000258665.

Moroz, L.L. 2014. The genealogy of genealogy of neurons. Commun Integr Biol 7(6):e993269. doi: 10.4161/19420889.2014.993269.

Moroz, L. L. 2015. Convergent evolution of neural systems in ctenophores. J Exp Biol 218, 598–611. doi: 10.1242/jeb.110692

Moroz, L.L. 2021. Multiple origins of neurons from secretory cells. Front Cell Dev Biol ;9:669087; doi: 10.3389/fcell.2021.669087.

Moroz, L.L. 2023. Syncytial nets vs. chemical signaling: emerging properties of alternative integrative systems. Front Cell Dev Biol;11:1320209; doi: 10.3389/fcell.2023.1320209.

Moroz, L.L. 2024. Brief history of ctenophora. Methods Mol Biol. ;2757:1–26. doi: 10.1007/978-1-0716-3642-8_1.

Moroz, L.L. (2026). Ctenophore neural systems: Independent origin(s) and “alien neurons”. In: Kaas, J.H., Herculano-Houzel, S. (eds.) Evolution of Nervous Systems, Third Edition. vol. 1, pp. 37–77. US: Elsevier doi.org/10.1016/B978-0-443-27380-3.00044-0

Moroz, L.L., Collins, R., Paulay, G. 2024 Ctenophora: Illustrated Guide and Taxonomy. Methods Mol Biol.,,,,,2757:27–102. doi: 10.1007/978-1-0716-3642-8_2.

Moroz, L. L., Kocot, K. M., Citarella, M. R., Dosung, S., Norekian, T. P., Povolotskaya, I. S., Grigorenko, A. P., Dailey, C., Berezikov, E., Buckley, K. M., Ptitsyn, A., Reshetov, D., Mukherjee, K., Moroz, T. P., Bobkova, Y., Yu, F., Kapitonov, V. V., Jurka, J., Bobkov, Y. V., Swore, J. J., Girardo, D. O., Fodor, A., Gusev, F., Sanford, R., Bruders, R., Kittler, E., Mills, C. E., Rast, J. P., Derelle, R., Solovyev, V. V., Kondrashov, F. A., Swalla, B. J., Sweedler, J. V., Rogaev, E. I., Halanych, K. M., Kohn, A. B. 2014. The ctenophore genome and the evolutionary origins of neural systems. Nature 510, 109–114. doi: 10.1038/nature13400

Moroz, L.L., Kohn, A.B. 2015. Unbiased View of Synaptic and Neuronal Gene Complement in Ctenophores: Are There Pan-neuronal and Pan-synaptic Genes across Metazoa? Integr Comp Biol 55, 1028–1049. doi: 10.1093/icb/icv104.

Moroz, L. L., Kohn, A. B., 2016. Independent origins of neurons and synapses: insights from ctenophores. Philos Trans R Soc Lond B Biol Sci 371, 20150041. doi: 10.1098/rstb.2015.0041.

Moroz, L.L., Nikitin, N.A., Poličar, P.G., Andrea B. Kohn A.B., Romanova, D.Y. 2021a. Evolution of glutamatergic signaling and synapses. Neuropharmacology;199:108740. doi: 10.1016/j.neuropharm.2021.108740.

Moroz, L. L., Romanova, D. Y., Kohn, A. B. 2021b. Neural versus alternative integrative systems: molecular insights into origins of neurotransmitters. Philos Trans R Soc Lond B Biol Sci 376, 20190762. doi: 10.1098/rstb.2019.0762.

Moroz, L. L., Sohn, D., Romanova, D. Y., Kohn, A. B., 2020. Microchemical identification of enantiomers in early-branching animals: Lineage-specific diversification in the usage of D-glutamate and D-aspartate. Biochem Biophys Res Commun 527, 947–952. doi: 10.1016/j.bbrc.2020.04.135.

Norekian, T.P., Moroz, L.L. 2016, Development of neuromuscular organization in the ctenophore Pleurobrachia bachei. J Comp Neurol524(1):136–151 doi: 10.1002/cne.23830.

Norekian, T.P., Moroz, L.L. 2019a. Neural system and receptor diversity in the ctenophore Beroe abyssicola. J Comp Neurol; 527(12):1986–2008; doi: 10.1002/cne.24633.

Norekian, T.P., Moroz, L.L. 2019b. Neuromuscular organization of the ctenophore Pleurobrachia bachei. J Comp Neurol.,,,,, 527(2):406–436; doi: 10.1002/cne.24546.

Norekian, T.P., Moroz, L.L. 2020. Comparative neuroanatomy of ctenophores: Neural and muscular systems in Euplokamis dunlapae and related species. J Comp. Neurol.,,,,, 528(3):481–501; doi: 10.1002/cne.24770.

Norekian, T.P., Moroz, L.L. 2021, Development of the nervous system in the early hatching larvae of the ctenophore Mnemiopsis leidyi. J Morphol 282(10):1466–1477 doi: 10.1002/jmor.21398.

Norekian TP, Moroz, L.L. 2024. Recording cilia activity in ctenophores. Methods Mol Biol. 2024;2757:307–313. doi: 10.1007/978-1-0716-3642-8_14.

Regnard, C., Desbruyeres, E., Huet, J. C., Beauvallet, C., Pernollet, J. C., Edde, B., 2000. Polyglutamylation of nucleosome assembly proteins. J Biol Chem 275, 15969–15976.

Sachkova MY, Nordmann EL, Soto-Àngel JJ, Meeda Y, Górski B, Naumann B, Dondorp D, Chatzigeorgiou M, Kittelmann M, Burkhardt P. 2021. Neuropeptide repertoire and 3D anatomy of the ctenophore nervous system. Curr Biol. 31(23):5274–5285.e6. doi: 10.1016/j.cub.2021.09.005.

Tamm, S. L. 2014. Cilia and the life of ctenophores. Invertebr. Biol.133, 1–46. doi: 10.1111/ivb.12042

Yang, W.-T., Hong, S.-R., He, K., Ling, K., Shaiv, K., Hu, J., Lin, Y.-C., 2021. The Emerging Roles of Axonemal Glutamylation in Regulation of Cilia Architecture and Functions. Frontiers in Cell and Developmental Biology 9.

Young, V. R., Ajami, A. M., 2000. Glutamate: an amino acid of particular distinction. J Nutr 130, 892S–900S.

Yu, A., Alberstein, R., Thomas, A., Zimmet, A., Grey, R., Mayer, M.L., Lau, A.Y., 2016. Molecular lock regulates binding of glycine to a primitive NMDA receptor. Proc. Natl. Acad. Sci. U. S. A. 113, E6786–E6795.

Vargas AM, DeBiasse MB, Dykes LL, Edgar A, Hayes TD, Groso DJ, Babonis LS, Martindale MQ, Ryan JF. 2024. Morphological and dietary changes encoded in the genome of Beroe ovata, a ctenophore-eating ctenophore. NAR Genom Bioinform. 6(2):lqae072. doi: 10.1093/nargab/lqae072.

Wang, W., Jack, B. M., Wang, H. H., Kavanaugh, M. A., Maser, R. L., Tran, P. V., 2021. Intraflagellar Transport Proteins as Regulators of Primary Cilia Length. Frontiers in Cell and Developmental Biology 9.

Wehland J, Willingham MC, Sandoval IV. 1983. A rat monoclonal antibody reacting specifically with the tyrosylated form of alpha-tubulin. I. Biochemical characterization, effects on microtubule polymerization in vitro, and microtubule polymerization and organization in vivo. J Cell Biol. Nov;97(5 Pt 1):1467–75. doi: 10.1083/jcb.97.5.1467.

Wehland J, Willingham MC. 1983. A rat monoclonal antibody reacting specifically with the tyrosylated form of alpha-tubulin. II. Effects on cell movement, organization of microtubules, and intermediate filaments, and arrangement of Golgi elements. J Cell Biol. Nov;97(5 Pt 1):1476–90. doi: 10.1083/jcb.97.5.1476.

Wulf E, Deboben A, Bautz FA, Faulstich H, Wieland T. 1979. Fluorescent phallotoxin, a tool for the visualization of cellular actin. Proc Natl Acad Sci U S A. Sep;76(9):4498–502. doi: 10.1073/pnas.76.9.4498.

